# Seasonal occurrence of Japanese pygmy octopus *Octopus parvus* in the intertidal zone

**DOI:** 10.1101/2021.03.14.435358

**Authors:** Yuta Yamate, Takumi Ohya, Toshifumi Wada, Takeshi Takegaki

## Abstract

The Japanese pygmy octopus *Octopus parvus* is a small octopus that occurs commonly along the coast of southern Japan, and is caught using traditional fishing methods. To date, however, there have been no studies on the ecology of this species. In this study, we investigated the annual seasonal occurrence of *O. parvus* in the intertidal zone of Fukue Island, Nagasaki, Japan, and examined the growth, maturation, and feeding habits of this octopus. We accordingly found that the octopus inhabits the intertidal zone of the study site from August to January, during which time it appears to move from the high to low intertidal zone, and subsequently migrates to the subtidal zone. During low tide at night in the intertidal zone, we observed that the octopuses actively captured and fed on prey, such as fish, crabs, hermit crabs, shrimp, snails, and bivalves. Both males and females were found to be characterized by similar body sizes and growth, although gonadal development occurs relatively earlier in males. However, any mating or brooding behaviors were not observed during the course of the study. Our observations thus indicate that *O. parvus* uses intertidal zones as feeding grounds for rapid growth and maturation, and may thereafter move to the subtidal zone for spawning.

## Introduction

The life span of most octopuses is typically less than a year (Mangold 1983; Boyle & Rodhouse 2008), during which time they grow large, mature, and reproduce. Octopuses are generally carnivorous animals that feed on a diverse range of organisms, such as crabs, shrimps, hermit crabs, snails, bivalves, fish, and even other cephalopods (Leite et al. 2009a; Hanlon and Messenger 2018). Rapid growth and maturation can be achieved by having access to abundant food resources (Giménez and García 2002; Semmens et al. 2004; André et al. 2009), and accordingly, an abundance of food may be an important factor determining habitat selection. Such rapid high growth would be presumably be advantageous in terms of both survival and reproduction.

Many species of octopus inhabit shallow coastal areas, and several are known to use intertidal zones (Iribarne 1991; Scheel 2002; Oosthuizen and Smale 2003; Leite et al. 2009a, b; Storero et al. 2010, 2013; Herwig et al. 2012; de Beer and Potts 2013; Wada 2017). Given their high productivity and biodiversity, intertidal zones represent an important ecosystem (Leigh et al. 1987; Underwood 2000) that would support an abundance of potential prey organisms for octopuses. For example, *Octopus tehuelchus* actively feeds and grows large in the intertidal zone during the warm months (Storero et al. 2010, 2013), whereas *Octopus insularis* uses the intertidal zone as a post-settlement nursery ground (Leite et al. 2009b). However, intertidal zones also tend to be unstable habitats in which environmental conditions, such as temperature, salinity, water levels, and dissolved oxygen levels, undergo marked diurnal and seasonal fluctuations (Truchot and Duhamel-Jouve 1980; Raffaelli and Hawkins 1996; Horn et al 1999; Helmuth and Hofmann 2001). In this regard, the daily activity patterns of intertidal octopuses are affected by tidal levels (Mather 1991; Roper and Hochberg 1988). As the temperature dropped at the beginning of fall, *O. tehuelchus* females move from the intertidal to subtidal zone to spawn and tend eggs (Iribarne 1991). Furthermore, in some species, mating behavior has been observed in the intertidal zone during low tide, both at night (Wada 2017) and during the daytime (Huffard 2007; Huffard et al. 2008).

The Japanese pygmy octopus *parvus* is a comparatively small octopus (10–15 cm in total length) endemic to Japan that inhabits coastal areas south of the Boso Peninsula and Sagami Bay (Sasaki 1929; Kubodera 2013). This species is caught in the intertidal zone from summer to fall using traditional fishing methods, which entail throwing ash or salt into rock holes during periods low tide (Sauer et al. 2019). Currently, however, there is relatively little information available regarding the temporal and functional usage of the intertidal zone by this species. Although females of *O. parvus* tending eggs in nests have been observed in the intertidal zone during spring (Takegaki and Wada, unpublished data), there has of yet been no reports regarding the reproductive season and behavior of this species. In this study, we investigated the occurrence of *O. parvus* in the intertidal zone throughout the year and examined the species’ growth, maturation, and feeding habits.

## Materials and methods

### Distribution survey and specimen collection

Field investigations were conducted once monthly from September 2018 to August 2019 (with the exception of February 2019) in the rocky intertidal zone along the Odomari coast of Fukue Island, Nagasaki Prefecture, Japan (32°38ʹ45″N, 128°48ʹ44″E), wherein numerous shallow intertidal pools (40 cm in maximum depth) are formed the during periods of low tide. The water temperature in the mid-tide zone ranges from 8 to 35°C. Given that *O. parvus* individuals typically emerge from their nests during the night, we conducted field surveys for 2 to 3 h during the night-time low tide periods of spring tides. We walked in groups of two to five persons in the intertidal zone with a flashlight (600 lm), searching for octopuses by turning over rocks in tidepools, and collecting specimens using a hand net and large tweezers. As our surveys of tidepools were performed as a group, the number of investigators would not appear to have a significant effect on the number of octopuses observed. To obtain a larger number of specimens, the survey area was extended to the western and lower intertidal zones after October 2018 (ca. 260 m × 90 m).

To investigate seasonal change in the occurrence of these octopuses, we recorded the longitudinal and latitudinal coordinates of each collection point using digital cameras (Tough TG-5; Olympus, Tokyo, Japan, WG-4 GPS; Ricoh, Tokyo, Japan) or a smartphone (iPhone 7; Apple Inc., California, USA) equipped with a GPS function (accuracy: <10 m), even in cases in which we failed to capture the octopus (37 of 140 observed individuals; Table 1). Using the coordinate data thus obtained, we plotted the geographical locations of octopus sitings on a map of the study area. In certain instances, coordinate data obtained in September (*n* = 1) and October (*n* = 8) in 2018 were not used, as they showed spurious point data (e.g., points located offshore or in inland areas), which could probably be attributed to a temporary failure to detect any satellite signals.

**Table 1.**
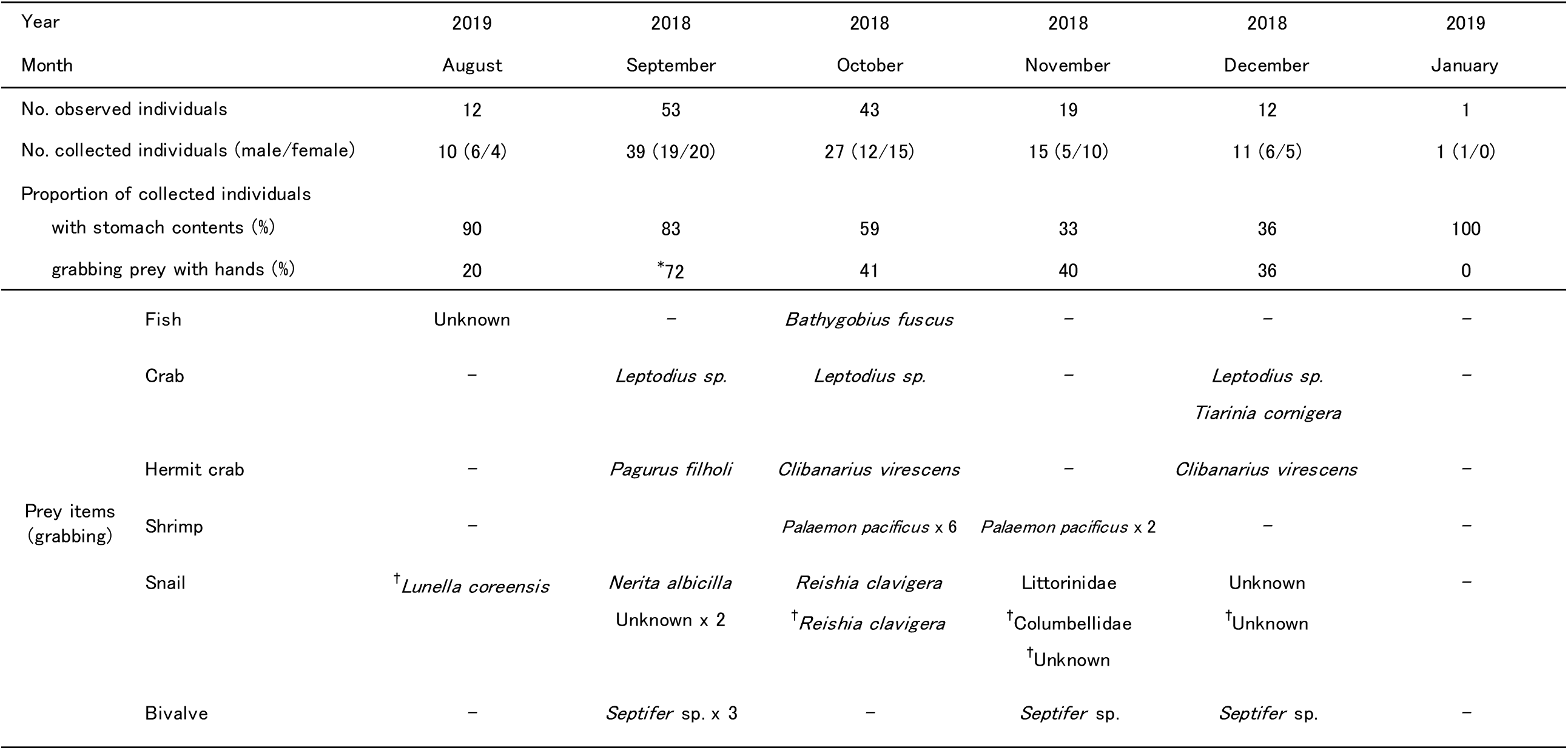
The numbers of observed and collected samples for each month. Monthly change in the proportion of individuals with stomach contents and individuals grabbing prey with hands, and prey items that octopus was grabbing at the time of collection in the field (numbers represent sample size). *: note that the proportion was calculated on the basis of 18 samples (details in the text). †: empty snail shell (either snail or hermit crab).

If, at the time of collection, octopuses were observed capturing prey organisms, these animals were also collected to characterize the feeding habits of *O. parvus*. However, a few prey samples obtained from 21 individuals collected in September 2018 were lost in the field. The octopuses thus collected were anesthetized with 1% ethanol prior to storing at -30°C for subsequent analysis.

### Measurement of specimens

After thawing, the ventral mantle length (VML) of octopus specimens was measured using a caliper (to the nearest 0.05 mm) as an index of body size, and wet body weight was measured using an electronic microscale (0.1 g: CP124S; Sartorius AG, Göttingen, Germany). To evaluate the degree of maturation based on a gonadosomatic index (GSI: gonad weight/body weight × 100), gonads were dissected and weighed (0.001 g). For measurement purposes, weighed male gonads included the testis, vas deferens, spermatophoric gland, spermatophore storage, and accessory spermatophoric gland, whereas the female gonads included the ovary, oviductal gland, proximal oviduct, and distal oviduct (González-Gómez et al. 2018). The presence of stomach content was confirmed, and where possible, the prey collected concomitantly with octopuses was identified to the species level.

### Data analysis

To examine the effects of season (month) and sex on the body size (VML), generalized linear models (GLMs) with Gaussian distribution and log-link function were used. The monthly changes in GSI values were analyzed using Tukey HSD multiple comparison test. All analyses were conducted using R Studio ver. 3. 6. 2 (R Core Team 2019).

### Results

*Octopus parvus* was observed in the intertidal zone of the study site from September 2018 to January 2019 and in August 2019 (Fig. 1), with the number of individuals peaking in September (*n* = 53) and October (*n* = 43), and subsequently undergoing a gradual decline until January of the following year (*n* = 1). No individuals, including newly settled juveniles, were observed between March and July 2019. In October, octopuses were observed throughout the surveyed area from high to low intertidal zones, whereas from November to January, there was a decline in the number of individuals detected in the high intertidal zone, with the majority of observations being confined to the low intertidal zone (Fig. 1). There was not much difference in monthly occurrence and movement between males and females (Table 1, Fig.1). Although multiple individuals were sometimes found inhabiting the same tidepool, during the entire course of the field survey, any reproductive behaviors, such as courtship displays, copulation, or male–male competition over females, or indeed any other interactions among individuals, were not observed.

**Fig. 1.**
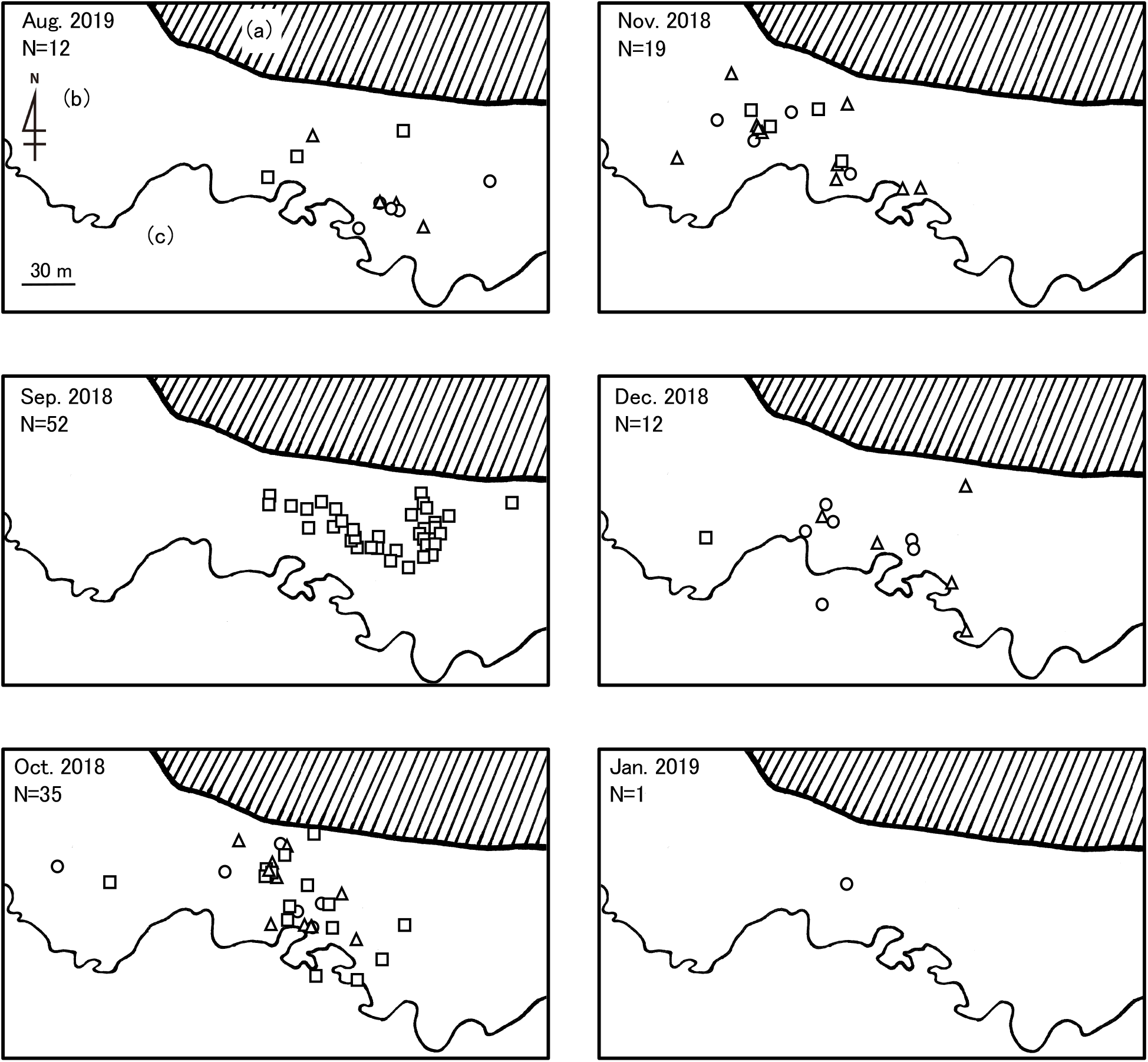
Seasonal changes in the location of *Octopus parvus* observations (open circles). (a) supralittoral zone (shaded area); (b) intertidal zone; (c) subtidal zone. Open circles and triangles indicate males and females, respectively; open squares indicate individuals of unknown sex. Note that several open circles overlap at the same site. Furthermore, we did no survey the western coastal area or lower intertidal zone in September 2018 (please refer to details in the main text).

The body size of collected individuals significantly increased from summer to winter, particularly between the months of August and October (41% and 61% increases in the mean size of males and females, respectively: GLMs, *χ*^*2*^ = 37.29, *p* < 0.0001; Fig. 2, Table 2). There was no significant effect of sex on body size (GLMs, *χ* ^*2*^ = 0.506, *p* = 0.477; Fig. 2, Table 2). The GSI of males significantly increased from September to November (Tukey HSD test, September to October: *p* < 0.01; October to November: *p* < 0.05), and was maintained at a high value throughout December and January (Fig. 3), whereas for females, there was a slightly prolonged significant increase in GSI from October to December (Oct to Nov and Nov to Dec, both *p* < 0.01; Fig. 3).

**Table 2.**
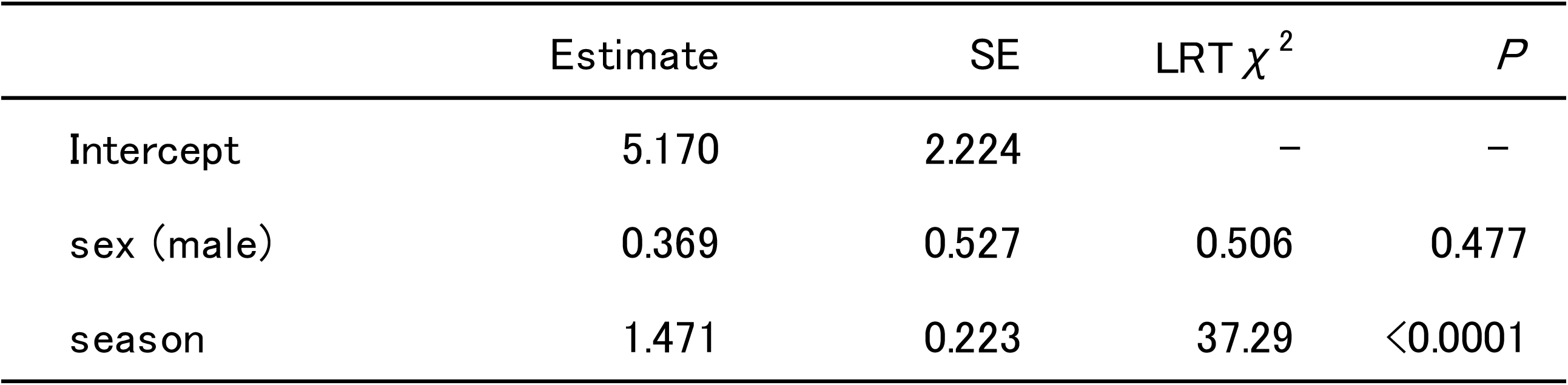
Effects of the season (month) and sex on the body size (VML) of octopus.

| | Estimate | SE | LRT $\chi^2$ | *P* |
| --- | --- | --- | --- | --- |
| Intercept | 5.170 | 2.224 | – | – |
| sex (male) | 0.369 | 0.527 | 0.506 | 0.477 |
| season | 1.471 | 0.223 | 37.29 | <0.0001 |

**Fig. 2.**
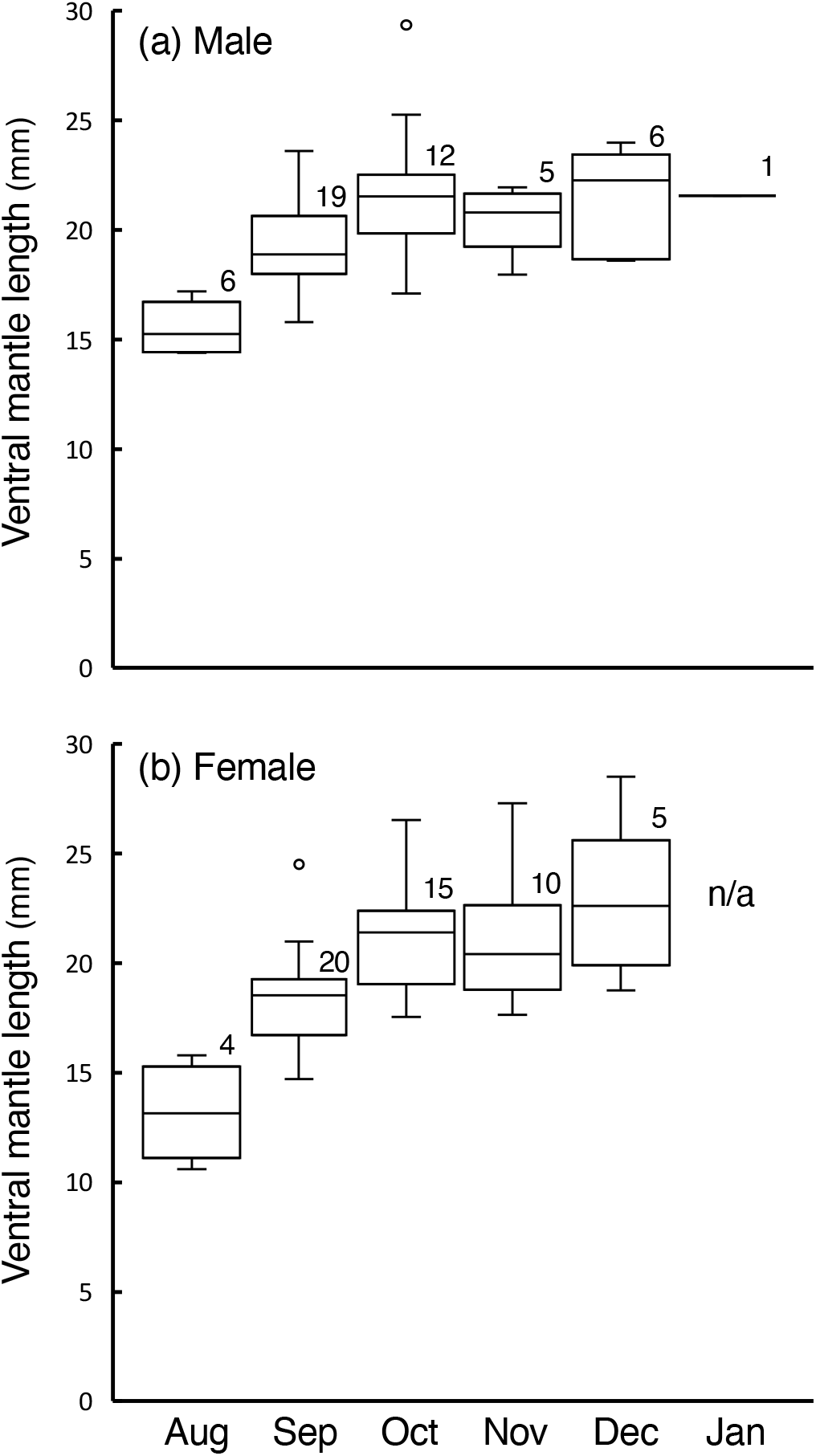
Seasonal changes in the ventral mantle length of (a) male and (b) female *Octopus parvus*. The numbers indicate the sample sizes for each month. The boxplots show medians, 25% and 75% quartiles, 10% and 90% percentiles (whiskers), and outliers (dots). Note that August and January specimens were collected in 2019, and others were collected in 2018.

**Fig. 3.**
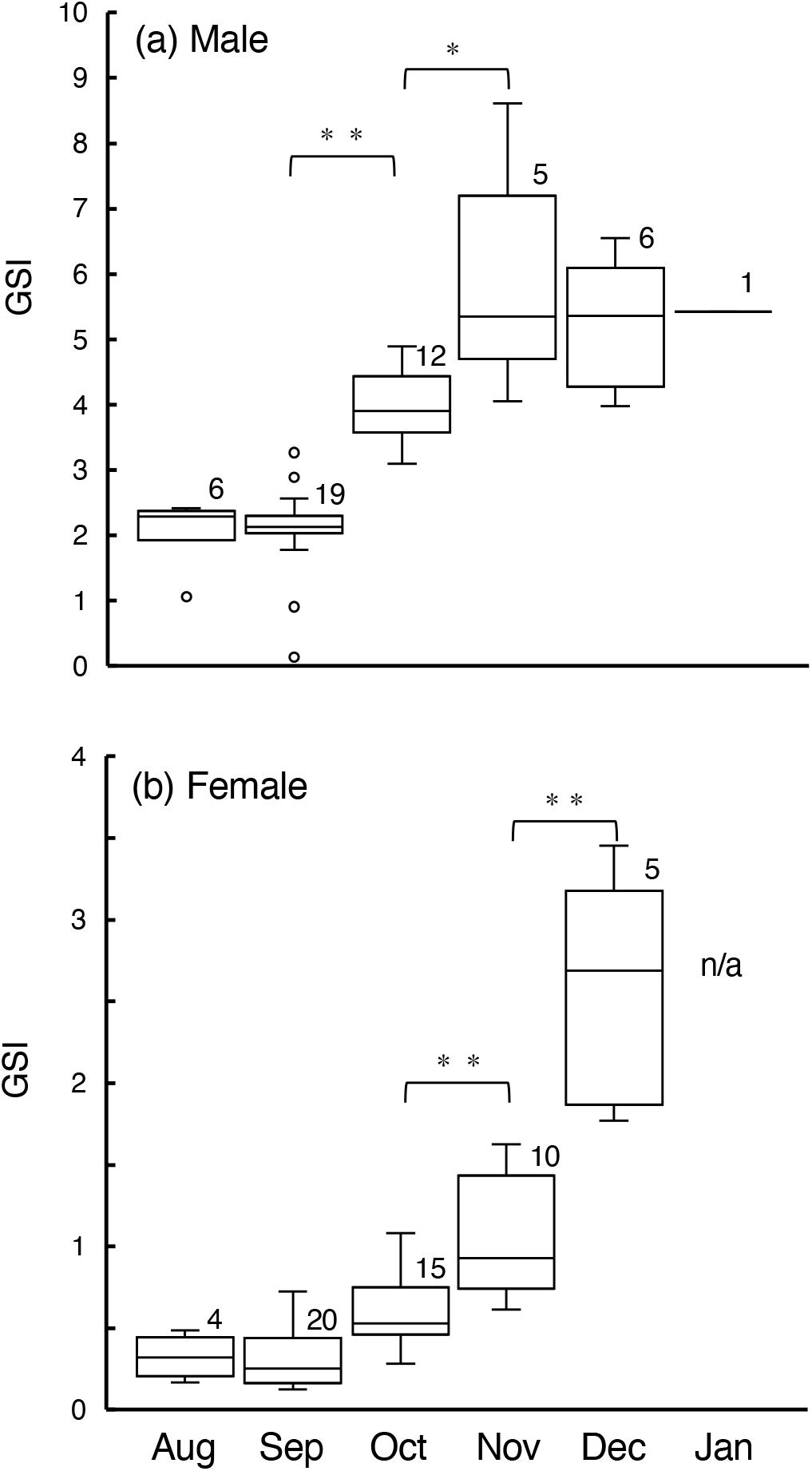
Seasonal changes in the gonadosomatic index (GSI) of (a) male and (b) female *Octopus parvus*. The numbers indicate the sample sizes for each month. The boxplots show medians, 25% and 75% quartiles, 10% and 90% percentiles (whiskers), and outliers (dots). Note that August and January specimens were collected in 2019, and the others were collected in 2018. *: P < 0.05, **: P < 0.01.

On a number of occasions, we observed octopuses capturing and consuming prey (fish, crabs, hermit crabs, shrimp, snails, and bivalves) at the time of collection (Table 1), and among the individuals collected in August and September, a high percentage had prey contents in the stomach, although the number declined gradually thereafter (Table 1).

## Discussion

Field investigations conducted throughout the year revealed that the Japanese pygmy octopus *parvus* uses the intertidal zone from August to January at this study site. Contrastingly, we observed no *O. parvus* individuals at the site between March and July, and we suspect that there were probably few, if any, in February. Monthly changes in the locations at which octopuses were observed tended to indicate that this species moves from the high to low intertidal zone during summer and winter months, and subsequently moves to the subtidal zone. Given that the temperature of water in the intertidal zone can undergo marked fluctuations in response to changes in air temperature, we speculate that *O. parvus* may move to the subtidal zone where the water temperature is higher and more stable during the colder months. In this regard, it has been suggested that in *O. tehuelchus*, a species showing a similar pattern of seasonal movement from intertidal to subtidal zones, low temperatures contribute to reducing female fecundity (Iribarne 1991).

As clearly indicated by daytime fishing activities, *O. parvus* tends to remain in rock holes during the periods of daytime low tides, whereas during low tide at night, individuals typically emerge from these crevices and actively hunt and consume on prey. Indeed, from August to December, the period during which *O. parvus* inhabits the intertidal zone, average body weight increased approximately 2.8-fold, thereby indicating that these octopuses use the intertidal zone as feeding grounds for rapid growth. A majority of the crustaceans, bivalves, and gastropods inhabiting in the rocky intertidal zone are potential prey items for octopus (Hanlon and Messenger 2018), and given that the *O. parvus* individuals collected in this study were confirmed to have fed on a diverse range of prey species, as predators, they may conceivably have a notable impact on the intertidal ecosystem. Conversely, we found no evidence to suggest predation or predatory attack against *O. parvus*, nor did we observe the presence of any potential predators such as large carnivorous fish or cephalopods. Accordingly, it is probably that use of the intertidal zone by *O. parvus*, may contribute to reducing risk of predation during this period, when small octopuses would more vulnerable to predatory attack (Katsanevakis and Verriopoulos 2004; Leite et al. 2009b).

During the months when octopuses were observed in the intertidal zone, both males and females were found to have similar body size and show comparable growth, and body sizes tended to be relatively constant after October, at which time individuals probably attained approximately maximal size, at least among males. However, gonad development was observed to occur relatively earlier in males, which is consistent with the earlier maturation of males reported in a number of other octopus species (*O. tehuelchus*: Iribarne 1991; *O. vulgaris*: Rodríguez-Rúa et al. 2005; *O. pallidus*: Leporati et al. 2008; and *O. maya*: Avila-Poveda et al. 2009; Markaida et al. 2016), and is conceivably associated with female acceptance of copulation prior to the maturation of ovaries and retention of sperm in the oviductal gland until spawning (González et al. 2011; de Lima et al. 2014; Hanlon and Messenger 2018). Although we did not conduct histological analyses in the present study, on the basis of the observed changes in GSI values, it is assumed that the males of *O. parvus* are reproductively viable in December, at the latest. Moreover, whereas nocturnal mating behavior has previously been observed at low tide in a similar small intertidal octopus *Abdopus* sp. (Wada 2017), any reproductive behaviors of *O. parvus* were not observed in the present study. However, the assumed seasonal migration of *O. parvus* from intertidal to subtidal zones is consistent with the autumnal movements of *O. tehuelchus* females, with the density of brooding females subsequently being observed to be higher in the subtidal zone (Iribarne 1991). In this regard, it is assumed that the mating of *O. tehuelchus* occurs immediately prior to the movement to the subtidal zone (Iribarne 1991). Accordingly, if mating in *O. parvus* precedes their migration to the subtidal zone, we speculate that it may take place during periods that were not monitored in present study, for example, during periods of neap or high tides. Nevertheless, given the observed decline in the number of individuals in the intertidal zone at the time of peak male GSI, it is conceivable that mating does not occur until reaching the subtidal zone.

Although we found no individuals at the study site during the period from March to July, two *O. parvus* females tending eggs in underwater nests were confirmed in April 2016 in the intertidal zone along the Miezaki coast, Nagasaki, at a site located 90 km distant from Fukue Island (Takegaki and Wada, unpublished data). Despite several daytime searches at low tide in April and May for nesting and egg-tending *O. parvus* females and eggs in rock holes, any females or eggs were not observed (Yamate and Takegaki, unpublished data). In this regard, we can speculate that if spawning and the tending of eggs do occur in the intertidal zone, there is a high probability that the females would be caught by fishermen in spring and summer. Thus, female spawning and egg care may be performed mainly in the subtidal zone. Furthermore, we were unable to confirm the recruitment of young individuals in this study, and given that adults initially appeared in the intertidal zone in August, it is conceivable that young individuals move from subtidal to intertidal zones after growth.

This study demonstrated the seasonal occurrence of the Japanese pygmy octopus *O. parvus* in the intertidal zone, which we assume the species uses as a feeding ground for rapid growth and maturation. Furthermore, they move from high to low intertidal zones during the summer and winter months, and probably subsequently migrate to the subtidal zone. They may act as an important key predator affecting the intertidal ecosystem, both spatially and temporally. However, to gain a more complete understanding of the life history of *O. parvus*, particularly with respect to recruitment and reproduction, it will be necessary to conduct further investigations, notably focusing on their use of the subtidal zone.

## Acknowledgements

We would like to thank Mr. T. Tanaka for providing information regarding *Octopus parvus* in the waters off Goto Island, and the Fisheries Cooperative Association of Oh-hama, Goto Fukue, for permission to carry out our field survey and collect specimens.

## Author contributions

Conceptualization, Y.Y., T.W. and T. T.; Methodology, Y.Y., T.W. and T. T.; Investigation, Y.Y., T.O., T.W. and T. T.; Writing – Original Draft, Y.Y. and T. T.; Writing – Review & Editing, Y.Y. and T. T.; Project Administration and Funding Acquisition, Y.Y., T.W. and T. T.

## Compliance with ethical standards

### Ethical approval

This study was approved by the Animal Care and Use Committee of the Faculty of Fisheries, Nagasaki University (permission no. NF-0021), in accordance with the Guidelines for Animal Experimentation of the Faculty of Fisheries (fish, amphibians, invertebrates), and Regulations of the Animal Care and Use Committee, Nagasaki University.

### Informed consent

Informed consent was obtained from all individual participants included in the study.

